# Resting bilateral sensorimotor mu rhythm suppression facilitates ipsilesional M1 excitability after stroke

**DOI:** 10.64898/2026.08.07.743250

**Authors:** Uttara U Khatri, Tharan Suresh, Joshua R Tatz, Sara J Hussain

## Abstract

**Objective:** Stroke-related corticospinal tract (CST) disruption causes lasting hand impairments, but many stroke survivors retain some residual CST connections. In neurotypical adults, motor cortex (M1) TMS interventions can strengthen CST transmission when coupled to EEG brain states reflecting heightened M1 excitability. Because stroke alters the relationship between these brain states and cortical excitability, we aimed to identify poststroke brain states that accurately capture ipsilesional M1 excitability. We hypothesized that heightened ipsilesional M1 excitability would be represented by a common, group-level EEG pattern and a participant-specific, personalized pattern.

**Methods:** We acquired single-pulse TMS-EEG-EMG datasets in 15 chronic stroke survivors with residual CST connections. We then identified group-level and individual-specific EEG power patterns that distinguished between high and low ipsilesional M1 excitability states.

**Results:** At the group level, bilateral sensorimotor mu power was significantly suppressed during high versus low excitability states, but this suppression did not correlate with hand impairment severity or trait-level ipsilesional M1 excitability. At the individual level, spatiotemporally varied EEG activity patterns distinguished between excitability states, but these patterns were only present in 60% of individuals.

**Conclusion and Significance:** This study is the first to systematically characterize poststroke EEG brain states reflecting ipsilesional M1 excitability. Findings suggest that individual-specific EEG patterns may inconsistently index ipsilesional M1 excitability and instead identify bilateral sensorimotor mu power suppression as a group-level excitability marker that is present across the full spectrum of poststroke hand impairment.

**Highlights:**

- We analyzed TMS-EEG-EMG to identify group and individual level ipsilesional motor cortical excitability states in chronic stroke
- Bilateral sensorimotor mu suppression marked heightened ipsilesional motor cortical excitability across hand impairment severity
- 60% participants had individual level scalp patterns linked to motor cortical excitability states, challenging their reliability

## Introduction

The corticospinal tract (CST) is the primary descending pathway supporting skilled hand control in humans (Lemon, 2008; Lemon & Griffiths, 2005; Welniarz et al., 2017). Stroke-related CST damage is thus a major cause of hand impairment, but ∼85% of stroke survivors have residual CST connections (Stinear et al., 2012, 2017). The presence and integrity of these remaining connections is a strong predictor of upper extremity recovery potential after stroke (Stinear, 2010; Zhu et al., 2010). Strengthening these residual CST connections could therefore enhance skilled control of paretic hand muscles, improve engagement with rehabilitation activities, and potentiate poststroke hand recovery.

Transcranial magnetic stimulation (TMS) offers one noninvasive means of strengthening residual CST connections (Di Lazzaro et al., 2008; Du et al., 2016; Jo & Perez, 2020; Smith & Stinear, 2016; Urbin et al., 2021; Ziemann & Siebner, 2015). Early work performed in small samples of neurotypical individuals showed that motor cortex (M1) TMS interventions increase CST transmission by inducing long-term potentiation (LTP)-like plasticity within motor cortical circuits that synapse with the CST (Chen et al., 1997; Kobayashi et al., 2004; Pascual-leone et al., 1994; Rizzo et al., 2004). However, more recent studies performed in larger samples showed that these same interventions do not reliably increase CST transmission (Hamada et al., 2013; López-Alonso et al., 2014; Schilberg et al., 2017). That is, M1 TMS interventions produce highly variable outcomes both within and across participants (Hussain & Freedberg, 2025). However, such TMS interventions are typically uncoupled from endogenous fluctuations in M1 excitability, which likely contributes to their weak and variable LTP-like effects (Bergmann et al., 2016; Buzsáki & Draguhn, 2004; Goldsworthy et al., 2014; Haegens et al., 2011; Hussain & Freedberg, 2025; Thut et al., 2017; Ziemann & Siebner, 2015; Zrenner et al., 2016, 2018).

Brain state-dependent TMS can overcome this limitation by delivering individual TMS pulses or pulse trains during electroencephalography (EEG)-defined brain states reflecting increased M1 excitability. Such brain states are defined using motor-evoked potential (MEP) amplitudes (Zrenner et al., 2018) and are often characterized by sensorimotor oscillatory power, phase, or their interaction. In neurotypical adults, M1 excitability positively correlates with sensorimotor mu (8-13 Hz; Karabanov et al., 2021; Thies et al., 2018) and beta (13-35 Hz) rhythm power (Hussain, Claudino, et al., 2019; Hussain, Cohen, et al., 2019). M1 excitability also varies significantly across mu and beta phases (Hussain, Claudino, et al., 2019; Ozdemir et al., 2022; Wischnewski et al., 2022; Zrenner et al., 2018), such that motor-evoked potentials (MEPs) are largest during mu trough and beta peak phases. Moreover, brain state-dependent TMS interventions targeting high-excitability mu trough phases increase CST transmission and improve motor learning, while identical interventions delivered outside of these phases do not (Hussain et al., 2021; Zrenner et al., 2018). As a whole, these findings strongly suggest that poststroke TMS interventions may only potentiate residual CST transmission and improve paretic hand control when coupled to brain states reflecting increased ipsilesional M1 excitability.

To date, all poststroke brain state-dependent TMS interventions have been delivered during mu rhythm trough phases (Lieb et al., 2023; Mahmoud et al., 2024). Because stroke causes adaptive and maladaptive changes within distributed sensorimotor cortical networks and descending systems (Grefkes & Ward, 2014; Larivière et al., 2018; Mooney et al., 2023; Sahrizan et al., 2025; Sato et al., 2022; Xu et al., 2015), these same brain states may not accurately index ipsilesional M1 excitability after stroke. Consistent with this possibility, recent work suggests that the effect of mu rhythm phase on M1 excitability is attenuated in stroke survivors with more severe upper extremity impairments (Brancaccio et al., 2025; Wischnewski et al., 2025). Further, mu trough-coupled rTMS does not improve motor recovery more than conventional brain-state independent rTMS in chronic stroke survivors with residual CST connections (Mahmoud et al., 2024). These findings suggest that poststroke rTMS must be delivered during alternative brain states that accurately capture ipsilesional M1 excitability in individuals with a wide range of motor impairments. However, such states have yet to be identified.

A major barrier to identifying brain states reflecting heightened ipsilesional M1 excitability is the high inter-individual variability in lesion characteristics, CST damage, and recovery trajectory among stroke survivors (Grefkes & Ward, 2014; Larivière et al., 2018; Stinear et al., 2017). However, this wide inter-individual variability is superimposed upon the human CST’s known neuroanatomical organization. Based on this rationale, we hypothesized that EEG patterns reflecting increased ipsilesional M1 excitability would be represented by both a common, group-level activity pattern and a participant-specific, personalized pattern. To test our hypothesis, we analyzed resting EEG and EMG recordings obtained during single-pulse ipsilesional M1 TMS in 15 chronic stroke survivors with residual CST connections. Group-level analyses revealed that bilateral sensorimotor mu rhythm power suppression reflects increased ipsilesional M1 excitability. In contrast, individual-specific analyses primarily showed that beta power suppression also reflected increased M1 excitability, although the scalp distribution of this pattern varied across individuals. Further, personalized EEG activity patterns could only be detected in ∼60% of individuals. Overall, our findings establish sensorimotor mu desynchronization as a physiologically informed, group-level target for poststroke brain state-dependent rTMS while also suggesting that personalized EEG activity patterns may not consistently index ipsilesional M1 excitability after stroke.

## Methods

### Data acquisition

#### Participants

Fifteen chronic stroke survivors (5 males, 10 females; age = 63 ± 2.3 years) who had experienced a stroke at least 6 months ago participated in a two-session study that included clinical evaluation, TMS, EEG, and EMG recordings. Session 1 involved clinical motor assessments and screening for residual CST connections to any paretic upper extremity muscle. Session 2 involved single-pulse TMS to the ipsilesional M1 during simultaneous EEG and EMG recordings. This study was performed at two sites: the University of Iowa, and the University of Texas at Austin. All procedures were approved by the Institutional Review Boards of each site, and all participants provided their written informed consent prior to participation.

#### Eligibility

Eligibility was evaluated on Day 1 using a combination of clinical assessments and TMS screening for residual CST connections. Inclusion criteria included ischemic or hemorrhagic stroke onset ≥ 6 months prior to enrollment, age ≥ 18 years, ability to provide informed consent, and the presence of detectable upper extremity hemiparesis. Exclusion criteria included any prescription medication changes within the past month, presence of other major neurological disorders (i.e., Parkinson’s disease, multiple sclerosis), and presence of TMS contraindications, score < 24 on the Mini-Mental State Examination, or inability to understand the study’s purpose and procedures. Participants were considered to have detectable upper extremity hemiparesis if they met one or more of the following criteria: score < 66 on the upper extremity Fugl-Meyer Assessment, score < 70 on a modified version of the Wolf Motor Function Test (all items except the key and lock task), ≥ 10% longer time to complete the 9-Hole Peg Test using the affected than the unaffected hand, ≥ 10% lower force production during precision, key, or power grip using the affected than the unaffected hand. Precision, key and power grip measurements were collected using a handheld analog dynamometer.

Participants were considered to have residual CST connections if a reliable and discernible motor-evoked potential (MEP) could be recorded from any resting paretic upper extremity muscle following ipsilesional single-pulse TMS. If MEPs could be reliably elicited from multiple paretic upper extremity muscles, the most distal muscle in which MEPs were reliably observed was selected for future measurements, with preference given to the first dorsal interosseous (FDI). The scalp hotspot was defined as the site at which largest and most reliable MEPs could be discerned for the target muscle. Participants in whom paretic upper extremity MEPs could not be elicited were ineligible for the study. Overall, 32 stroke survivors enrolled in this study and 16 met our eligibility criteria. Of these individuals, 15 completed the study. See *Table 1* for a description of participants, lesion characteristics and target muscles.

**Table 1.** Participant characteristics. Clinical and demographic characteristics of 15 chronic stroke survivors who completed the study. FMA = Fugl-Meyer Upper Extremity Assessment (maximum = 66), WMFT=Wolf Motor Function Test (maximum = 70, item 15 [key and lock] not tested). N/A = MRI scans or radiological reports for these participants could not be obtained.

| ID | Lesioned hemisphere | Months post stroke | FMA score | Age | Sex | Target muscle | Lesion location | WMFT score |
| --- | --- | --- | --- | --- | --- | --- | --- | --- |
| 1 | Left | 27.3 | 65 | 53 | F | FDI | subcortical | 60 |
| 2 | Right | 15.03 | 40 | 68 | M | FDI | both | 54 |
| 3 | Right | 38.17 | 33 | 73 | M | FDI | both | 48 |
| 4 | Left | 43.67 | 56 | 69 | F | FDI | subcortical | 70 |
| 5 | Right | 9.27 | 47 | 52 | F | FDI | subcortical | 68 |
| 6 | Right | 26.63 | 43 | 46 | M | FDI | both | 46 |
| 7 | Right | 305.23 | 62 | 71 | F | FDI | both | 68 |
| 8 | Right | 6.90 | 64 | 58 | M | FDI | cortical | 66 |
| 9 | Left | 39.23 | 49 | 63 | F | FDI | subcortical | 46 |
| 10 | Right | 9.00 | 21 | 51 | F | APB | subcortical | 27 |
| 11 | Right | 9.00 | 66 | 74 | F | FDI | both | 69 |
| 12 | Right | N/A | 46 | 71 | M | FDI | N/A | N/A |
| 13 | Left | 11.33 | 64 | 66 | F | FDI | N/A | 67 |
| 14 | Right | 39.53 | 4 | 66 | M | ADM | N/A | 0 |
| 15 | Right | 15.76 | 10 | 62 | F | FDI | N/A | 14 |

#### EEG and EMG acquisition

62-channel EEG signals (10-10 system) were recorded at 5 kHz (low-pass hardware filter cutoff frequency: 1250 Hz; 0.001 µV resolution) using TMS-compatible amplifiers (NeurOne Tesla, Bittium Biosignals, Finland). Impedances were maintained below 10 kΩ. We also recorded bipolar EMG signals from the most distal paretic muscle in which MEPs could be reliably observed at 5 kHz (low-pass hardware filter cutoff frequency: 1250 Hz; 0.001 µV resolution) using Ag-AgCl adhesive electrodes arranged in a belly-tendon montage. EEG and EMG signals were acquired during single-pulse TMS delivery.

#### Transcranial magnetic stimulation

Participants deemed eligible on Day 1 returned on Day 2 for a session of single-pulse TMS, EEG, and EMG recordings. After EEG electrode preparation, we identified the scalp hotspot and RMT for the most distal paretic muscle in which MEPs could be reliably observed. Hereafter, we refer to this muscle as the target muscle. To maximize transsynaptic activation of CST neurons, single-pulse TMS was delivered using an air-cooled figure-of-eight coil held at ∼45° relative to the mid-sagittal line (Mills et al., 1992; Deymed Diagnostic, XT100, biphasic pulse shape). The hotspot was identified as the scalp site at which suprathreshold single-pulse TMS elicited the largest and most reliable MEPs, and when possible, a focal muscle twitch. We then determined resting motor threshold (RMT) using an adaptive threshold-tracking tool (MTAT 2.0; Awiszus, 2011). RMT was on average 67.6 ± 4.6% (range = 42-100%) of maximum stimulator output.

Next, we delivered 6 blocks of 100 single-pulse TMS pulses at 120% RMT (inter-stimulus interval: 3 seconds ± random jitter) while simultaneously recording EEG and EMG signals. In 2 participants, 120% of RMT exceeded maximum stimulator output, and stimulation was instead applied at 100% of maximum stimulator output. In these two individuals, 100% of stimulator output corresponded to 100% and 110% RMT. During stimulation, participants rested quietly with their eyes open. To maintain a consistent attention level and prevent drowsiness during TMS delivery, participants were instructed to mentally count the number of TMS pulses delivered during each block. Participants were asked to report their pulse count to the research team at two random time points per block. Short breaks (2-10 minutes) were provided between blocks. Throughout the experiment, coil position accuracy and precision were monitored online using frameless neuronavigation (BrainSight, Rogue Research, Inc.). See Figure 1A for a visual depiction of the experimental timeline.

**Figure 1.**
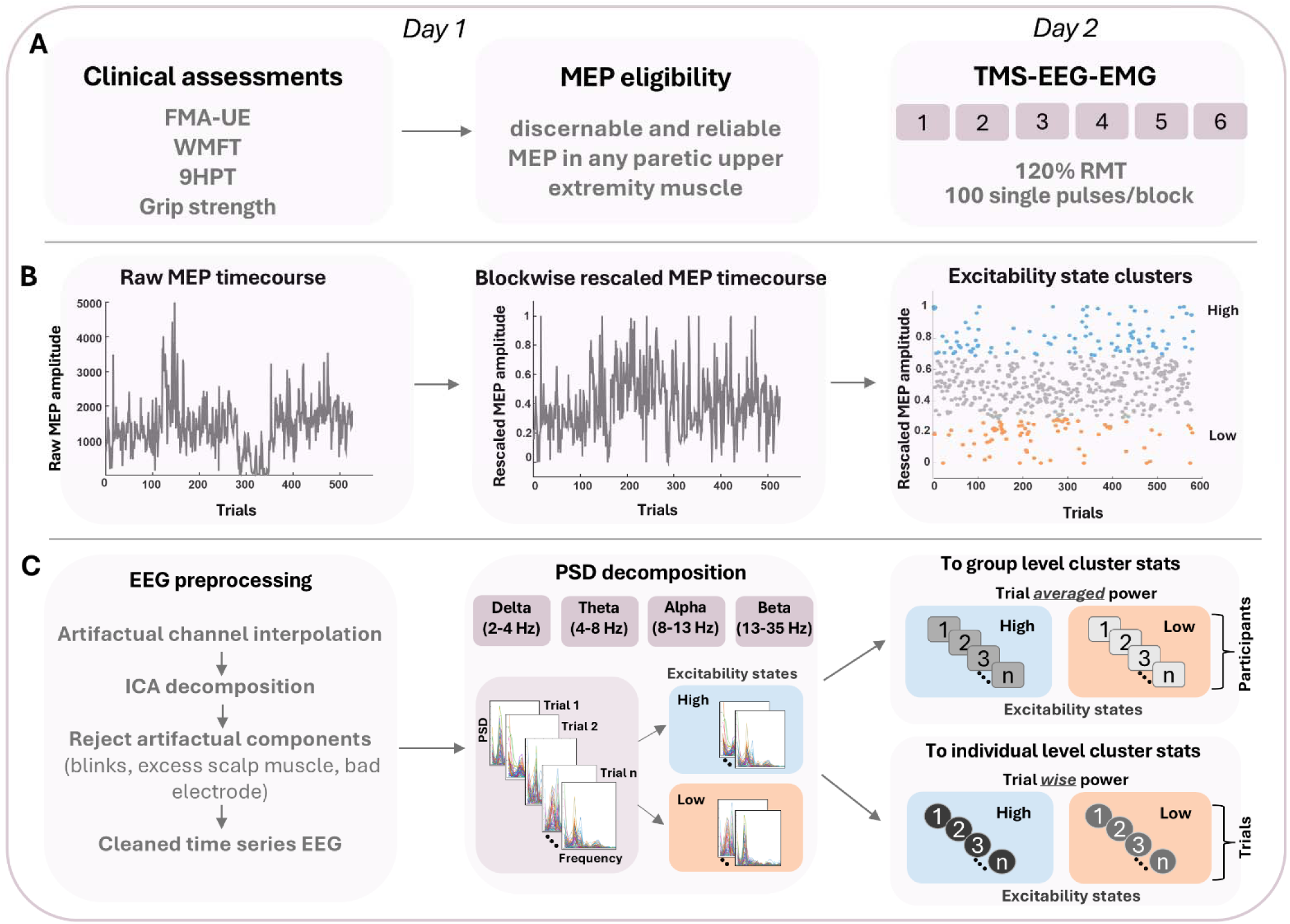
Depiction of experimental timeline and data analysis pipeline. A) Experimental timeline. B) Time course of paretic hand muscle MEP amplitudes obtained during 6 blocks of single-pulse ipsilesional M1 TMS. To capture trial-by-trial variability within each block rather than slow fluctuations in MEP amplitudes, MEP amplitudes obtained during each block were detrended, z-scored and rescaled from 0-1. Next, rescaled MEPs were clustered into high (blue circles) and low (orange circles) excitability states using k-means clustering (k = 4). C) Artifactual channels were interpolated and EEG artifacts were attenuated using independent components analysis. EEG data were then spectrally decomposed, and band-specific power was computed for delta, theta, alpha and beta bands for each trial. Resulting EEG power values were then binned into high and low excitability states per participant. Trial-averaged EEG power values obtained for each excitability state and frequency band were calculated, pooled, and forwarded to group-level statistical analysis. Trial-specific EEG power values obtained for each excitability state and frequency band were also calculated and forwarded to participant-specific statistical analysis.

### Data analysis

#### EEG preprocessing

EEG data were preprocessed using the EEGLAB toolbox (version 2024.1) and custom-written MATLAB (version R2025a) scripts. 62-channel EEG data were segmented from -2.005 to -0.005 seconds before each TMS pulse. Bad channels were identified using two criteria: (1) visual identification of channels exhibiting excessive 60 Hz line noise, high impedance, or movement artifacts (e.g., due to hand placement on the electrode), and (2) automated detection via kurtosis (capturing abnormal peakedness in the signal’s amplitude distribution, indicative of spike-like artifacts) and joint probability (capturing statistical improbability of a channel’s activity relative to other channels) measures calculated in EEGLAB. Channels exceeding seven standard deviations from the mean on either measure (computed across all channels within each block) were flagged as bad. Using these two approaches, 7.8 ± 0.6 channels were rejected per participant; these channels were subsequently interpolated using spherical spline interpolation across all blocks. EEG data were re-referenced to the common average and resulting pre-stimulus EEG data segments were demeaned and linearly detrended. Segments containing excessive scalp muscle activity, movement artifacts, and drift artifacts were discarded (14.2 ± 2.5 trials, 2.4 ± 0.4% of trials per participant). Remaining EEG segments were then low-pass filtered (cutoff = 50 Hz, zero-phase two-pass Hamming-windowed FIR filter, filter order = 1320). To account for rank reduction due to bad channel interpolation, ICA was performed using EEGLAB’s runica function with PCA rank adjustment enabled; this procedure matched the number of principal components to the effective data rank. Independent component analysis was used to identify components reflecting eye blinks, movement artifacts, scalp muscle activity, and cardiac activity via the Extended Infomax algorithm. Components consistent with these artifacts were manually identified and rejected (16.9 ± 0.8 independent components per participant). After rejection, remaining components were back-projected to the sensor space for further analysis (Figure 1C).

#### MEP analysis

Continuous EMG recordings obtained from the target muscle were segmented from -0.100 to +0.400 seconds relative to the TMS pulse, demeaned, and linearly detrended. We calculated MEP amplitudes as the peak-to-peak difference within a participant-specific post-stimulus time window. To adequately capture MEPs across all trials, this window was individually defined for each participant. Trials in which MEP responses could not be reliably detected from background EMG activity were excluded from further analyses (1.6 ± 0.7 trials, 0.2 ± 0.1% of trials per participant).

Next, we identified trials containing excessive pre-stimulus EMG activation. After attenuating line noise (discrete Fourier transform filter, frequencies removed = 30, 60, 90, 120, 150, 180 Hz), root-mean-square pre-stimulus EMG activation was calculated within a 75 ms window (0.100 to -0.025 seconds before each TMS pulse). Trials containing pre-stimulus EMG activation levels that exceeded two standard deviations above mean pre-stimulus activity were excluded from further analysis (20.2 ± 3.2 trials, 3.4 ± 0.5% of trials per participant).

Single-pulse TMS can produce slow drifts in MEP amplitudes (Pellicciari et al., 2016). The purpose of this study was to identify EEG activity patterns predicting trial-by-trial variation in ipsilesional M1 excitability rather than cumulative changes over time. To attenuate the influence of any cumulative excitability changes and instead capture trial-by-trial variation in MEP amplitudes, each block of MEP amplitudes was demeaned, linearly detrended, z-scored and rescaled from 0-1. Rescaled MEPs were then combined across blocks. For each participant, we then used k-means clustering (k = 4) of trial-by-trial rescaled MEP amplitudes to identify trials containing the largest and smallest MEPs. Trials belonging to the topmost and the bottommost clusters were retained for further analysis; hereafter, we refer to these trials as high and low excitability states (see Figure 1B). Overall, this approach identified 78.4 ± 15.4 trials corresponding to high excitability states and 159.4 ± 18.2 trials for low excitability states.

#### Hand impairment index

Hand impairment metrics are often highly correlated after stroke (Mirdamadi et al., 2023). We therefore calculated a composite hand impairment index for each stroke survivor by applying probabilistic principal components analysis to Fugl-Meyer hand/wrist subscores, 9-Hole Peg Test performance asymmetry, key grip force asymmetry, precision grip force asymmetry, and power grip force asymmetry. Fugl-Meyer hand/wrist subscores were expressed as one minus the proportion of the maximum possible score on the hand/wrist subscales (1 – [score/24]). For the 9-Hole Peg Test, we calculated the number of pegs placed and removed per second. Asymmetry scores for the 9-Hole Peg Test and all pinch/grip force assessments were calculated as (unaffected hand – affected hand)/(unaffected hand + affected hand). The first principal component explained 93.2% of total variance in all input metrics and was thus used as a composite hand impairment index (Mirdamadi et al., 2023). One participant did not complete all clinical motor assessments and was therefore excluded from hand impairment analyses.

#### Group-level cortical activity patterns

Preprocessed scalp EEG data were used to identify activity patterns that distinguished between high and low excitability states at the group level. First, the EEG channels for participants with left hemisphere stroke (N = 4) were mirrored across the midline, such that all EEG data reflected a right hemisphere lesion. Next, each participant’s pre-stimulus whole-scalp EEG data were spectrally decomposed between 2 and 35 Hz with 1 Hz frequency resolution (multi-taper method with Hanning windows). Then, power spectra obtained for each channel, frequency and trial were normalized to the mean spectrum observed for that channel and frequency across all trials. For each excitability state, we binned normalized spectra into canonical frequency bands (delta = 2-4 Hz, theta = 4-8 Hz, alpha = 8-13 Hz, and beta = 13-35 Hz) and averaged the resulting power values across trials. Cluster-based permutation tests were used to identify scalp channel clusters at which these EEG power values significantly differed between high and low excitability states.

#### Participant-specific, personalized cortical activity patterns

The same approach described above was used to identify participant-specific activity patterns that distinguished between high and low excitability states. Because all analytical procedures were performed at the individual participant level, we retained each participant’s normalized power spectra for all trials and channels. Participant-level, cluster-based permutation tests were then used to identify scalp channel clusters at which band-specific power values significantly differed between high and low excitability states for each participant. See Figure 1C for a visual depiction of analytical procedures.

### Statistical analyses

All statistical analyses were performed in RStudio and MATLAB 2025A. Alpha was set to 0.05 for all comparisons and data are expressed as mean ± SEM. Model fits for linear mixed effects models were visually examined using histograms of residuals and quantile-quantile plots, and the significance of fixed effects was determined using an F-test with Satterthwaite’s denominator degrees of freedom (*anova* function in lmerTest; Bates et al., 2015; Kuznetsova et al., 2017). For generalized linear mixed effects models and generalized linear models, model fits were visually examined using histograms of residuals and quantile-quantile plots, and the significance of fixed effects was determined using Wald t-tests (summary function in lmerTest; Bates et al., 2015; Kuznetsova et al., 2017). Post hoc pairwise comparisons were performed by estimating marginal means and were corrected for multiple comparisons using the Benjamini-Hochberg false discovery rate (FDR) procedure (emmeans package; Benjamini & Hochberg, 1995).

#### Cluster-based permutation testing

We identified EEG power differences between high and low excitability states using cluster-based permutation tests implemented in FieldTrip (Maris & Oostenveld, 2007; Oostenveld et al., 2011). Separate tests were conducted for each frequency band. We used a within-subjects design for group-level analysis (unit of observation: subject, independent variable: excitability state) and a between-trials design for participant-level analysis (unit of observation: trial, independent variable: excitability state). For both analysis types, we first computed a cluster-based test statistic quantifying the effect of excitability state by calculating a t-value at every channel (group-level tests = dependent samples t-statistic, participant-level tests = independent samples t-statistics). Channels with t-values ≤ 2.5^th^ or ≥ 97.5^th^ percentiles (reflecting a two-sided t-test) were selected as candidate members of that cluster. Selected channels were then clustered based on spatial adjacency. Cluster-level statistics were calculated as the sum of all t-values and the cluster with the largest t-value was selected for further analysis. Next, permutation tests were used to estimate the p-value corresponding to this observed test statistic value. Here, the reference distribution of the permutation test was approximated using the Monte-Carlo method. To achieve this, all observations from both excitability states were pooled together in a single dataset. Data were then randomly partitioned by shuffling the relationship between observations and excitability states. Next, the test statistic (i.e., the maximum of cluster-level t-values) was calculated from this random partition. This procedure was repeated 1500 times and was used to construct an empirical null distribution of test statistics. Finally, the Monte Carlo significance probability for a two-sided hypothesis test was calculated as the proportion of random partitions that resulted in a larger t-statistic than the observed t-statistic for the selected cluster.

#### Group-level EEG pattern analysis

We first identified group-level differences in scalp EEG power between high and low excitability states using cluster-based permutation tests. After identifying EEG channels at which band-specific power significantly differed between high and low excitability states, we then evaluated if these EEG power differences could be caused by differences in pre-stimulus EMG activation between states. This model focused only on the mean EEG power from significant channels identified during cluster-based permutation testing. Given that the EEG power values followed a right-tailed skewed distribution, a generalized linear mixed effect model was used (gamma distribution function with a log link function). Here, trial-by-trial mean EEG power was the response variable, STATE was the fixed effect, trialwise pre-stimulus EMG activation was a covariate, and PARTICIPANT was a random effect.

Next, we evaluated whether the identified group-level pattern generalizes across the full range of observed band-specific EEG power values and MEP amplitudes. We tested for a continuous relationship using a trial-by-trial linear mixed-effects model, with rescaled MEPs as the response variable, mean POWER across channels as a fixed effect, pre-stimulus EMG activation as a covariate, and PARTICIPANT as a random effect. This model focused only on the mean EEG power from significant channels identified during cluster-based permutation testing.

Finally, we examined relationships between the group-level EEG activity pattern identified during cluster-based permutation testing, composite hand impairment, and RMT. Here, RMT was used as a proxy for trait-level ipsilesional M1 excitability, with larger RMT values reflecting lower excitability. For each participant, we calculated the ratio between band-specific power values averaged over significant channels from the group level cluster. We first confirmed that hand impairment scores, RMT, and EEG power ratios adhered to a normal distribution using Shapiro-Wilk tests. Then, we regressed group-level EEG power ratios against composite hand impairment score and RMT using Pearson’s correlation.

#### Participant-specific, personalized EEG pattern analysis

We identified participant-specific differences in scalp EEG power between high and low excitability states using cluster-based permutation tests. For each participant, separate cluster-based tests were used for each frequency band. Next, we tested whether participant-specific, personalized patterns identified during cluster-based permutation testing could be attributed to differences in pre-stimulus EMG activation. To achieve this, we used separate trial-by-trial generalized linear models for each participant and frequency band. These models focused solely on mean EEG power at channels identified during cluster-based permutation testing. Band-specific EEG power values were included as the response variable, STATE as a fixed effect, pre-stimulus EMG activation as included a covariate, and PARTICIPANT as a random effect.

Personalized EEG activity patterns that distinguished between high and low excitability states could only be identified in a portion of tested individuals. We therefore also explored differences in composite hand impairment and RMT between participants in whom personalized patterns could and could not be identified. Given the limited sample size of each subgroup, we performed separate exploratory Wilcoxson rank-sum tests to compare differences in hand impairment scores and RMT between participants who exhibited individual patterns and those who did not.

#### Relative strength of group-level and personalized activity patterns

In participants with detectable individual patterns, we qualitatively compared the extent to which group-level and personalized patterns covaried with excitability state. To achieve this, we calculated the Cohen’s d effect size for each participant using EEG power values obtained from the group-level pattern and that individual’s personalized pattern. For personalized patterns, effect sizes were calculated for each frequency band. The independent samples Cohen’s d effect size was used to account for differing trial numbers between high and low excitability states.

## Results

### Group-level activity patterns

We used cluster-based permutation tests to identify whole-scalp EEG patterns that distinguished between high and low excitability states for each frequency band. This approach identified no significant differences in delta, theta, or beta power between high and low excitability states (delta: cluster-specific p > 0.16, theta and beta: no candidate clusters).

However, the same statistical approach identified significant differences in mu-alpha power between high and low excitability states (Figure 2A-D). These differences were present over bilateral sensorimotor cortical areas (cluster-specific p < 0.006, cluster channels: CP4, CP6, C4, C6, FT8, F8, FC6, FC4, FC2, FCz, FC1, FC3, F2, Fz, F1, C3, CP3, CP1), with sensorimotor mu power being significantly lower during high than low excitability states (Figure 2E).

**Figure 2.**
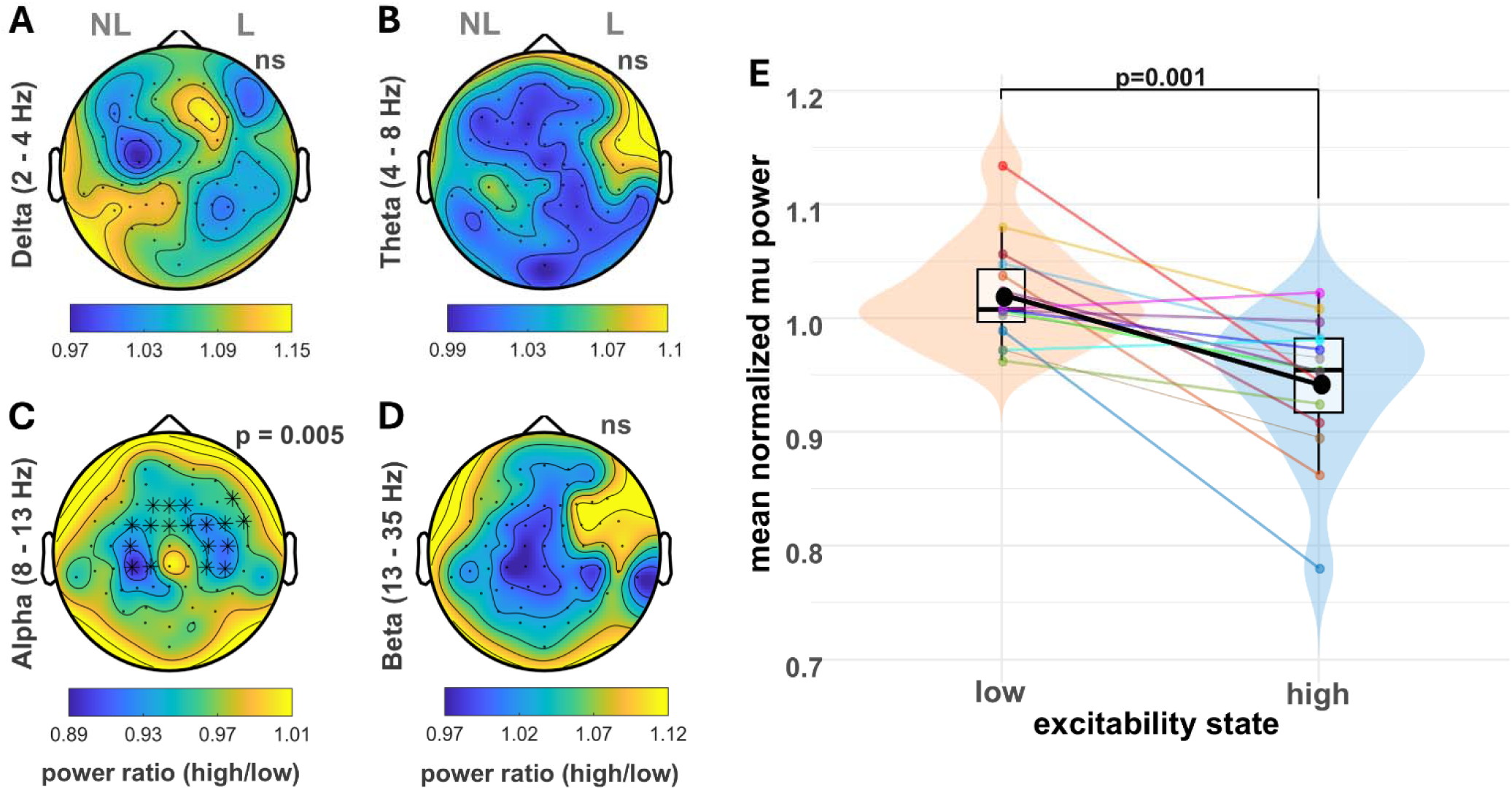
Group-level EEG patterns distinguishing between high and low ipsilesional M1 excitability states. A-D) Topoplots depicting the group averaged ratio between EEG power during high and low excitability states for each frequency band. Asterisks and p-values indicate significant differences between excitability states obtained using cluster-based permutation tests. E) Mean bilateral sensorimotor mu power during high and low excitability states. Colored circles represent individual participants, black circles represent group-level averages. Boxplots represent distribution for each group: the central line is the median, the box spans the interquartile range, and whiskers indicate the minimum and maximum observed values. The p- value indicates a significant difference in mean mu power obtained using group-level trial-by- trial linear mixed effects models. L=lesioned hemisphere, NL=non-lesioned hemisphere, ns=not significant.

Given that sensorimotor mu power suppression occurs during volitional movement (Pfurtscheller, 1997; Pfurtscheller and da Silva 1999), we next examined whether the difference in bilateral sensorimotor mu power between excitability states was driven by differences in pre- stimulus EMG activation. To achieve this, we used a trial-by-trial generalized linear mixed effects model that compared mean mu power between excitability states and included pre- stimulus EMG activation of the target muscle as a covariate. This analysis was restricted to significant channels identified during cluster-based permutation testing. We observed no significant relationship between pre-stimulus EMG activation and bilateral sensorimotor mu power (no effect of pre-stimulus EMG activation, t-value = -1.108, p = 0.267), and bilateral sensorimotor mu power differed significantly between excitability states even after including pre- stimulus EMG activation as a covariate (main effect of STATE, t-value = -3.180, p = 0.001).

Next, we tested for a continuous relationship between bilateral sensorimotor mu power and MEP amplitudes. This analysis focused on EEG and MEP data obtained throughout single-pulse TMS rather than only high and low excitability states. Here, we used a trial-by-trial linear mixed effects model that was restricted to significant channels identified during cluster-based permutation testing; this model also incorporated pre-stimulus EMG activation as a covariate. As expected, pre-stimulus EMG activation was significantly positively related to MEP amplitude (main effect of pre-stimulus EMG activation: F = 95.95, p < 0.0001). However, after accounting for trial-by-trial variation in pre-stimulus EMG activation, the relationship between bilateral sensorimotor mu power and MEP amplitude did not reach significance (no effect of mu power: F = 1.805, p = 0.179).

Finally, we evaluated relationships between the strength of the identified group-level bilateral mu power pattern, RMT, and a composite hand impairment score that integrated Fugl-Meyer hand/wrist subscores, 9-Hole Peg Test performance asymmetry, and precision, key, and power grip strength asymmetry. However, the ratio between bilateral sensorimotor mu power during high versus low excitability states did not correlate with composite hand impairment (r = -0.28, p > 0.3) or RMT (r = -0.01, p > 0.95). That is, brain state-dependent suppression of bilateral sensorimotor mu power did not covary with poststroke hand impairment or trait-level ipsilesional M1 excitability (see Figure 3A, 3B).

**Figure 3.**
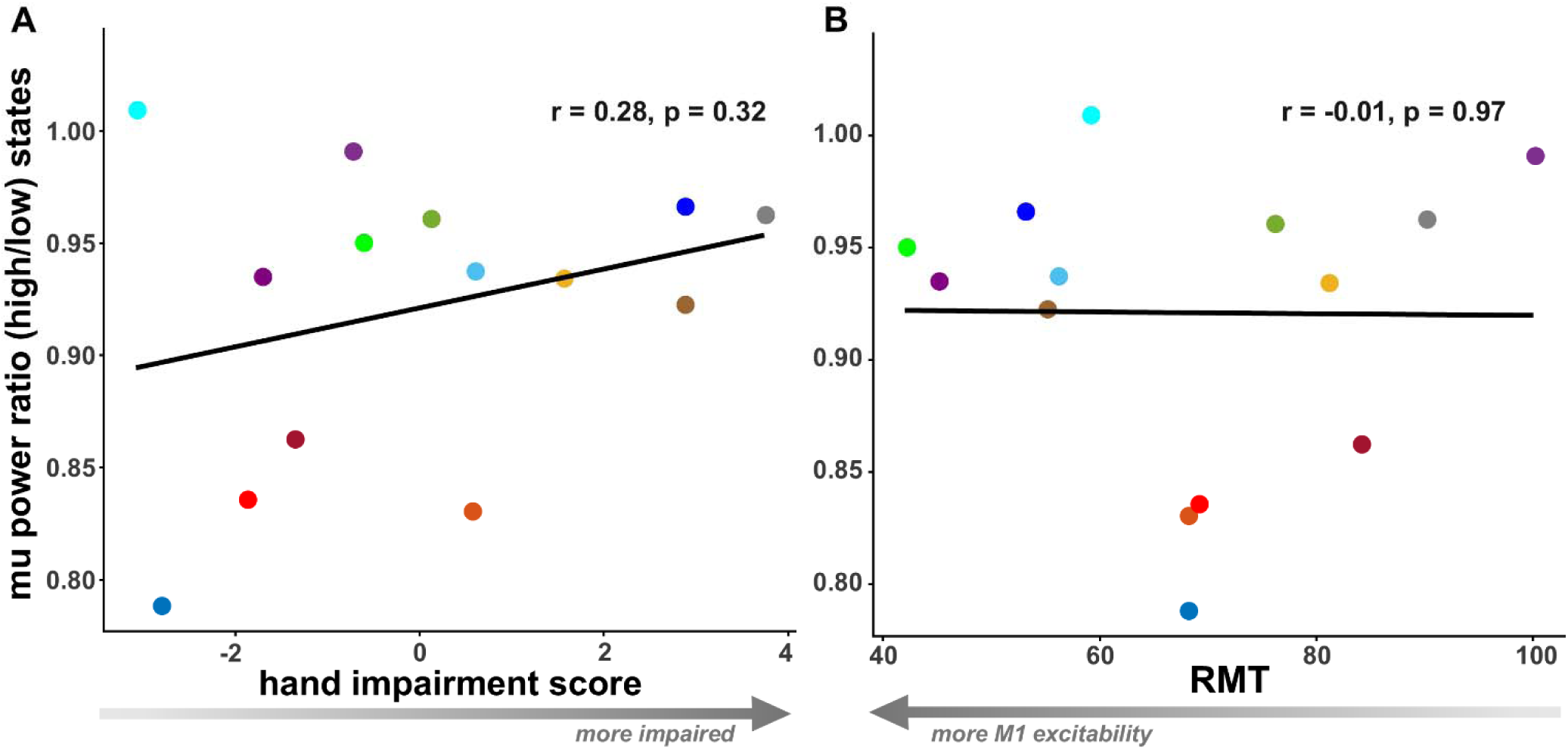
Relationships between group-level bilateral mu power suppression, hand impairment, and RMT. Relationships between the ratio of sensorimotor mu power between high and low excitability states and composite hand impairment (A) and RMT (B). For A, smaller/more negative hand impairment values reflect more severe impairment. For B, smaller RMT values reflect higher trait-level ipsilesional M1 excitability.

### Participant-level activity patterns

We also evaluated participant-specific differences in whole-scalp EEG activity using cluster-based permutation tests for each frequency band. These tests identified EEG power differences between excitability states in the theta, alpha, and beta ranges. See *Tables 2-4* for participant-level statistical results.

**Table 2.**
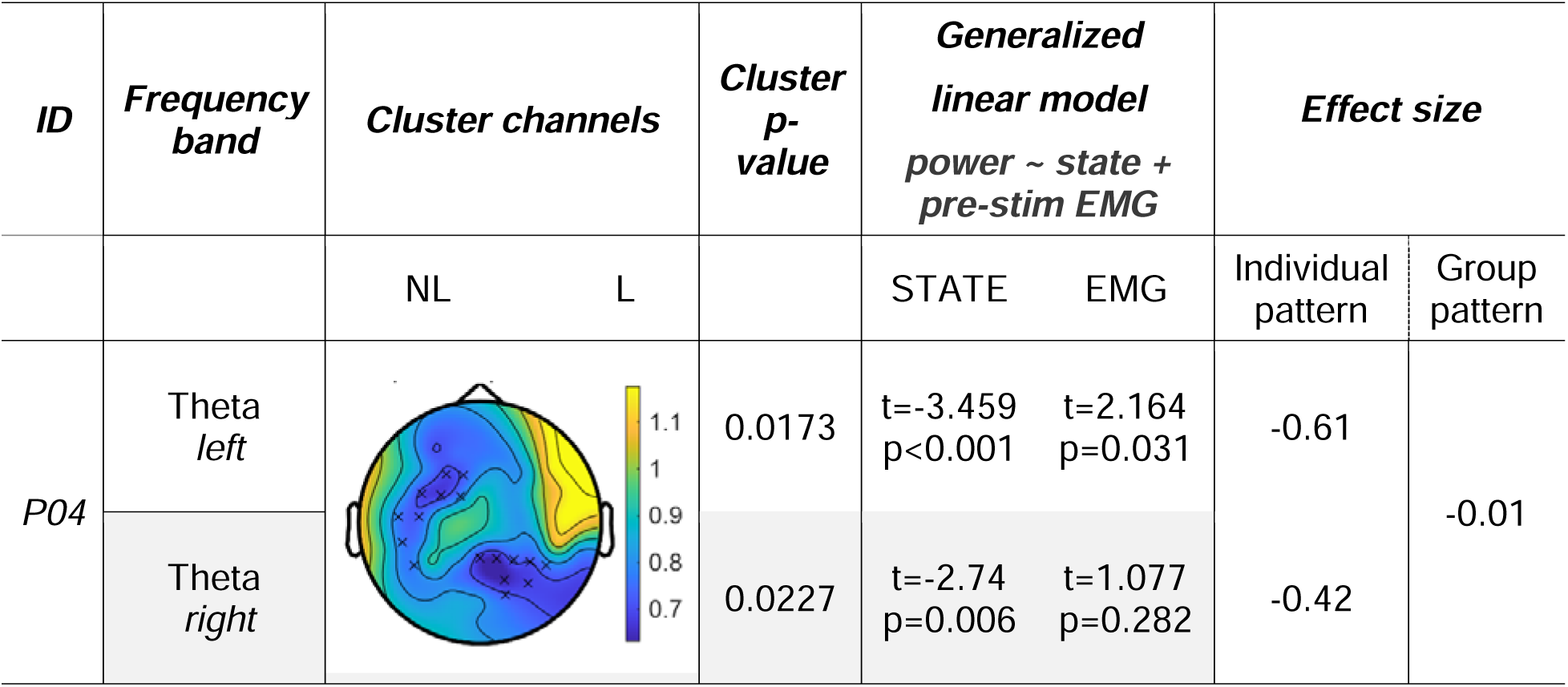
Theta EEG power patterns distinguishing between high and low excitability states in one participant. Topoplots depict the ratio of theta power between high and low excitability states. NL = non-lesioned hemisphere, L = lesioned hemisphere. Xs indicate p < 0.05.

| ID | Frequency band | Cluster channels |  | Cluster p-value | Generalized linear model<br>power ~ state + pre-stim EMG |  | Effect size |  |
| --- | --- | --- | --- | --- | --- | --- | --- | --- |
|  |  | NL | L |  | STATE | EMG | Individual pattern | Group pattern |
| P04 | Theta left |  |  | 0.0173 | t=-3.459<br>p<0.001 | t=2.164<br>p=0.031 | -0.61 | -0.01 |
|  | Theta right |  |  | 0.0227 | t=-2.74<br>p=0.006 | t=1.077<br>p=0.282 | -0.42 |  |

**Table 3.** Mu-alpha EEG power distinguishing between high and low excitability states in two participants. Topoplots depict the ratio of alpha power between high and low excitability states. NL = non-lesioned hemisphere, L = lesioned hemisphere. Xs indicate p < 0.05.

| ID | Frequency band | Cluster channels |  | Cluster p-value | Generalized linear model<br>power ~ state + pre-stim EMG |  | Effect size |  |
| --- | --- | --- | --- | --- | --- | --- | --- | --- |
|  |  | NL | L |  | STATE | EMG | Individual pattern | Group pattern |
| P10 | Alpha | | | 0.042 | $t=-3.330$<br>$p<0.001$ | $t=1.042$<br>$p=0.298$ | -0.42 | -0.07 |
| P11 | Alpha | | | 0.024 | $t=-3.019$<br>$p=0.003$ | $t=-0.964$<br>$p=0.335$ | -0.35 | -0.13 |

**Table 4.**
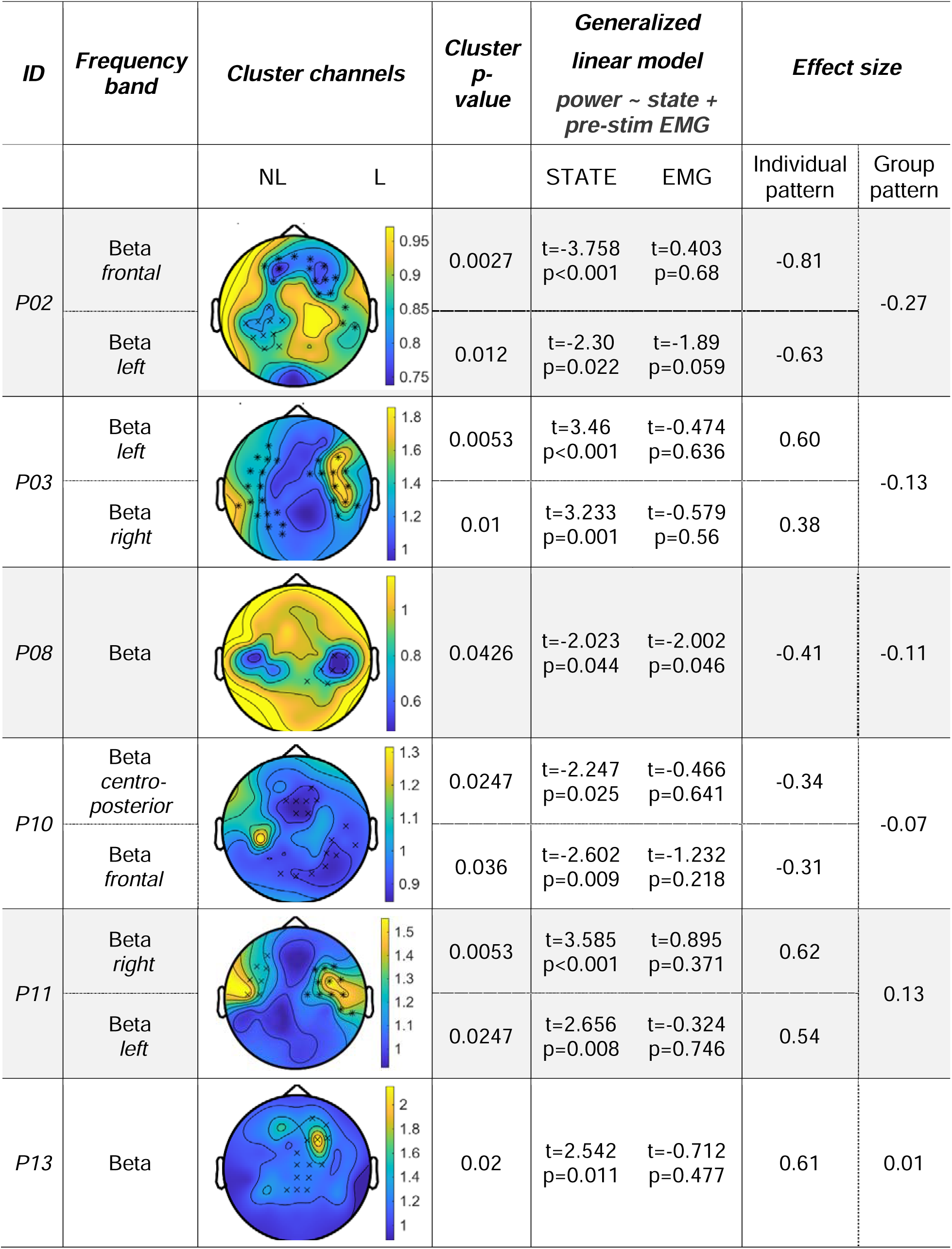
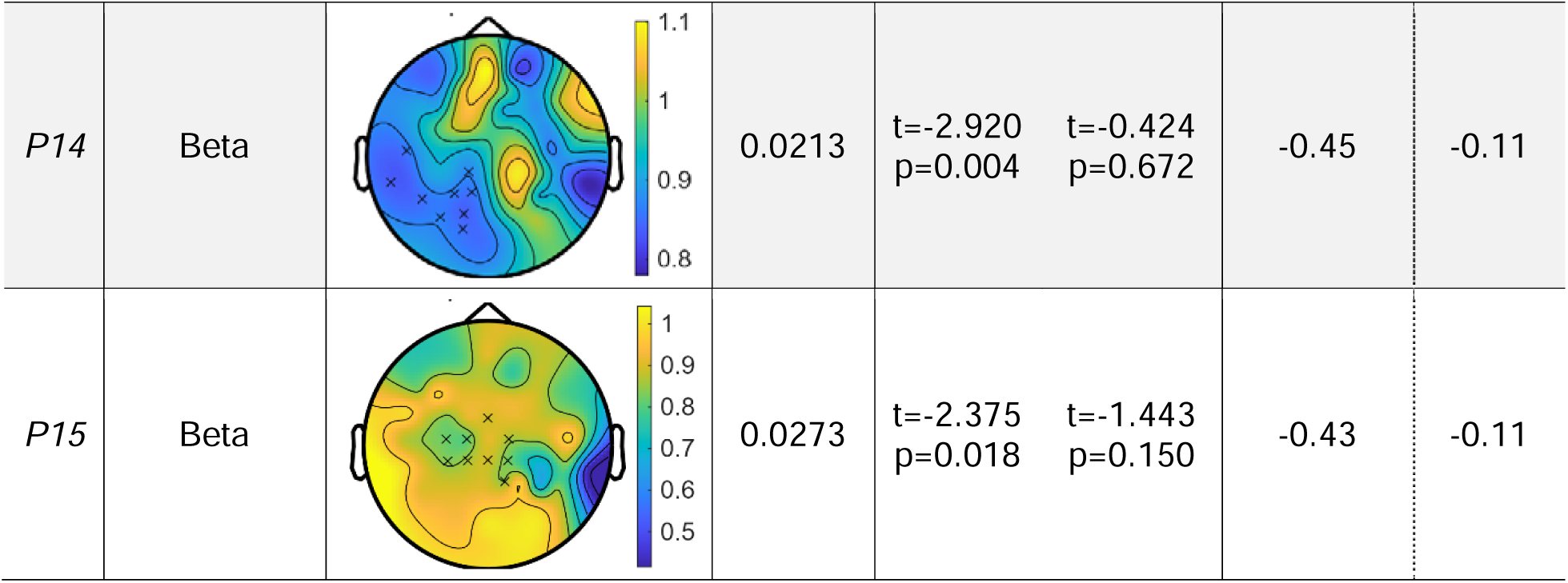
Beta EEG power distinguishing between high and low excitability states in eight participants. Topoplots depict the ratio of beta power between high and low excitability states. NL=non-lesioned hemisphere, L=lesioned hemisphere. Xs indicate p < 0.05 and asterisks indicate p < 0.001.

| ID | Frequency band | Cluster channels<br>NL L | Cluster p-value | Generalized linear model<br>power ~ state + pre-stim EMG |  | Effect size |  |
| --- | --- | --- | --- | --- | --- | --- | --- |
|  |  |  |  | STATE | EMG | Individual pattern | Group pattern |
| P02 | Beta frontal |  | 0.0027 | t=-3.758<br>p<0.001 | t=0.403<br>p=0.68 | -0.81 | -0.27 |
|  | Beta left |  | 0.012 | t=-2.30<br>p=0.022 | t=-1.89<br>p=0.059 | -0.63 |  |
| P03 | Beta left |  | 0.0053 | t=3.46<br>p<0.001 | t=-0.474<br>p=0.636 | 0.60 | -0.13 |
|  | Beta right |  | 0.01 | t=3.233<br>p=0.001 | t=-0.579<br>p=0.56 | 0.38 |  |
| P08 | Beta |  | 0.0426 | t=-2.023<br>p=0.044 | t=-2.002<br>p=0.046 | -0.41 | -0.11 |
| P10 | Beta centro-posterior |  | 0.0247 | t=-2.247<br>p=0.025 | t=-0.466<br>p=0.641 | -0.34 | -0.07 |
|  | Beta frontal |  | 0.036 | t=-2.602<br>p=0.009 | t=-1.232<br>p=0.218 | -0.31 |  |
| P11 | Beta right |  | 0.0053 | t=3.585<br>p<0.001 | t=0.895<br>p=0.371 | 0.62 | 0.13 |
|  | Beta left |  | 0.0247 | t=2.656<br>p=0.008 | t=-0.324<br>p=0.746 | 0.54 |  |
| P13 | Beta |  | 0.02 | t=2.542<br>p=0.011 | t=-0.712<br>p=0.477 | 0.61 | 0.01 |
| P14 | Beta |  | 0.0213 | t=-2.920<br>p=0.004 | t=-0.424<br>p=0.672 | -0.45 | -0.11 |
| P15 | Beta |  | 0.0273 | t=-2.375<br>p=0.018 | t=-1.443<br>p=0.150 | -0.43 | -0.11 |

Significant differences in theta power between high and low excitability states were present for only one participant (P04). In this individual, theta power was significantly weaker during high versus low excitability states at left fronto-central channels (cluster-specific p < 0.003) and at right parieto-occipital channels (cluster-specific p < 0.02).

Significant differences in alpha power between excitability states were present for two participants. In one individual (P10), alpha power was significantly lower during high versus low excitability states at right parieto-occipital channels (cluster-specific p < 0.05), while in the other individual (P11) power was significantly lower during high versus low excitability states at left centro-parieto-occipital channels (cluster-specific p < 0.03).

Significant differences in beta power between excitability states were present in eight participants. However, the scalp channels at which these differences were detected varied across individuals. In two participants, beta power over fronto-central areas differed significantly between high and low excitability states, with one participant showing weaker beta power during high versus low excitability states (P02) and the other participant showing stronger beta power during high versus low excitability states (P13; cluster-specific p < 0.04). Additionally, four participants showed significantly weaker centro-parieto-occipital beta power during high relative to low excitability states (P08, P10, P14, P15; cluster-specific p < 0.03). Additionally, two participants (P03, P11) showed higher beta band power at temporal channels during high versus low excitability states (cluster-specific p < 0.03).

Next, we tested whether differences in pre-stimulus EMG activation could explain the participant-level patterns identified via cluster-based permutation testing. To achieve this, we used separate trial-by-trial generalized linear mixed models that were restricted to significant channels identified during cluster-based permutation testing. These models also included pre-stimulus EMG activation of the target muscle as a covariate. We observed a significant relationship between pre-stimulus EMG activation and EEG power in two participants (significant effect of pre-stimulus EMG: p = 0.03 for P04; p = 0.046 for P08). However, EEG power differed significantly between excitability states even after accounting for pre-stimulus EMG activation in each participant (effect of STATE: p < 0.05 for all participants; see *Tables 2- 4*).

Overall, we identified personalized, participant-specific differences in EEG power between high and low excitability states in 9 of 15 participants, corresponding to 60% of our sample. We therefore explored whether composite hand impairment scores or trait-level ipsilesional M1 excitability systematically differed between participants in whom personalized patterns could (N = 9) and could not (N = 6) be identified. However, neither composite hand impairment scores nor RMTs differed between subgroups (p = 0.51 and p = 0.84; see Figure 4A and B). We also did not observe any qualitative differences in lesion location or hemisphere between participants in whom personalized patterns were and were not detected.

**Figure 4.**
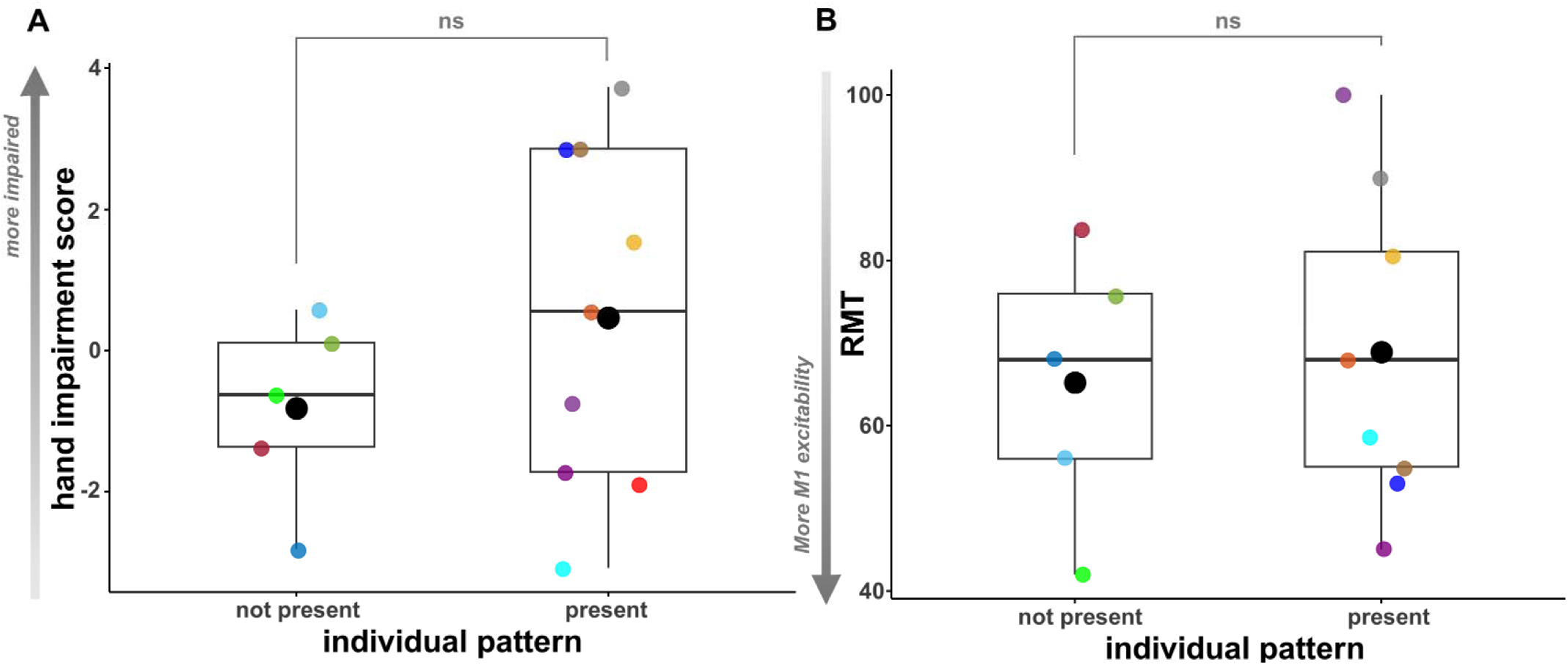
Differences in hand impairment and RMT between participants in whom participant-specific EEG patterns were and were not detected. Colored circles represent individual participants and black circles represent group-level averages. Boxplots represent distribution for each group: the central line is the median, the box spans the interquartile range, and whiskers indicate the minimum and maximum observed values.

Finally, we quantified the extent to which group-level and individual-specific EEG power patterns varied between high and low excitability states using Cohen’s d effect sizes. This analysis was only performed in individuals in whom a personalized EEG pattern could be detected (9 of 16 participants). For these participants, individual-specific EEG power differences showed an average effect size of 0.51 for the theta band (one participant, *Table 2*), 0.38 for the mu-alpha band (two participants, *Table 3*), and 0.51 for the beta band (eight participants, *Table 4*). In contrast, bilateral mu power differences identified during the group-level analysis showed an average effect size of 0.10 (*Tables 2-4*). Overall, the extent to which EEG power varied between high and low excitability states was stronger for individual-specific EEG patterns than for group-level patterns, but these personalized patterns were only detected in 60% of individuals.

## Discussion

In this study, we identified EEG brain states that distinguish between high and low ipsilesional M1 excitability in chronic stroke survivors with residual CST connections. We hypothesized that ipsilesional M1 excitability states would be captured by a common EEG activity pattern detectable at the group level and a personalized pattern detectable at the individual participant level. To test our hypothesis, we delivered single TMS pulses to the ipsilesional M1 during simultaneous EEG and EMG recordings. Group-level analyses revealed that bilateral sensorimotor mu rhythm power distinguished between excitability states, with mu power being significantly lower during high than low excitability states. Surprisingly, the magnitude of this mu power suppression did not depend on hand impairment severity. In contrast, participant-level analyses identified EEG power differences that differentiated between excitability states in 60% of tested participants; these differences were most common in the beta range. Although the direction, frequency, and scalp distribution of individual-specific EEG power differences varied across individuals, neither hand impairment severity nor trait-level ipsilesional M1 excitability could explain the presence or absence of such patterns. To our knowledge, this is the first study to systematically investigate and characterize EEG patterns distinguishing between high and low ipsilesional M1 excitability states after stroke.

Our main finding is that bilateral sensorimotor mu power was significantly suppressed during high relative to low excitability states. This finding has several implications for the design of poststroke brain state-dependent TMS interventions. First, our results broadly align with the established phenomenon of event-related desynchronization (ERD) during motor imagery, preparation, and execution (Pfurtscheller, 1997; Pfurtscheller & Aranibar, 1979; Stępień et al., 2011; Szurhaj et al., 2003; Takemi et al., 2013). The high excitability states characterized here may therefore reflect engagement of the same neural circuits driving voluntary movement, even at rest. Combined with the known role of mu ERD in volitional motor activity, our results suggest that these circuits may spontaneously cycle through windows of increased motor readiness, even when no overt movement is required. Second, poststroke brain state-dependent TMS interventions have primarily been coupled to ipsilesional mu rhythm trough phases (Lieb et al., 2023; Mahmoud et al., 2024). Although these phases reliably capture increased M1 excitability in neurotypical adults, this effect is only present when mu power is high (Hussain, Claudino, et al., 2019; Ozdemir et al., 2022; Suresh & Hussain, 2023). The suppressed mu power pattern identified in the current study is thus unlikely to co-occur with mu phase-dependent fluctuations in ipsilesional M1 excitability. Third, a recent study showed that the relationship between mu phase and ipsilesional M1 excitability is reduced in stroke survivors with greater upper extremity motor impairments (Wischnewski et al., 2025). However, in our study, the magnitude of mu power suppression between excitability states did not covary with hand impairment severity.

Fourth, mu trough phases reflect pulsed facilitation (Bergmann et al., 2019), while mu desynchronization captures GABA-ergic disinhibition (Takemi et al., 2013). Because cortical disinhibition is essential for motor cortex LTP (Hess & Donoghue, 1994), brain state-dependent TMS interventions targeting mu desynchronization could elicit stronger neuroplastic effects than those coupled to trough phases. As a whole, these observations suggest that mu power suppression and mu trough phases reflect physiologically distinct brain states and highlight the importance of considering each state’s properties when designing brain state-dependent TMS interventions.

Our findings should be interpreted in the context of prior studies evaluating relationships between mu power and M1 excitability in neurotypical adults. Sauseng et al. first reported that contralateral sensorimotor mu power recorded at rest negatively correlated with M1 excitability (Sauseng et al., 2009). Our results are consistent with these findings and expand them to the poststroke brain. However, Thies et al. later reported that sensorimotor mu power positively correlated with M1 excitability in neurotypical adults (Thies et al., 2018). Subtle methodological distinctions could feasibly account for these differences. For example, Thies et al. delivered EEG-informed TMS during pre-defined mu power levels detected from a single scalp channel, while Sauseng et al. and the current work used post-hoc trial sorting and whole-brain analyses to localize mu power differences between excitability states. While both studies reported relationships between local mu power and M1 excitability, the spatially diffuse pattern identified here is consistent with the notion that bilateral sensorimotor cortices co-activate following stroke (Schaechter et al., 2008; Stępień et al., 2011).

Participant-level analyses identified EEG patterns that distinguished between excitability states within theta, alpha, and beta bands in 9 of 15 participants. Of these patterns, beta power differed significantly between states in the majority of individuals (8 of 9), with beta power typically being suppressed during high relative to low excitability states. These findings align with the known phenomenon of movement-related sensorimotor beta power suppression (Barone & Rossiter, 2021; Pfurtscheller, 1997). However, three participants showed the opposite effect: beta power was significantly greater during high than low excitability states. In two of these individuals, beta power differences were present at temporal sites, which often show jaw muscle activity within the beta range (Whitham et al., 2007). Despite careful EEG artifact attenuation, we were unable to fully eliminate these likely artifacts. Regardless, participant-specific EEG patterns distinguishing between high and low excitability states most often reflected beta power suppression, although these patterns were too spatially variable to be observable at the group level. In contrast, none of the individual-specific EEG patterns identified here mimicked the group-level pattern (bilateral sensorimotor mu suppression). This discrepancy is likely due to limited signal-to-noise ratio of the group-level pattern, and future studies could feasibly strengthen it using volitional motor tasks (Pfurtscheller & Aranibar, 1979; Pfurtscheller & Lopes da Silva, 1999; Stępień et al., 2011; Takemi et al., 2013) or neurofeedback (Neuper & Pfurtscheller, 2010; Wang et al., 2019).

Surprisingly, participant-specific EEG patterns that distinguished between high and low excitability states were only identified in 60% of tested individuals. Although these patterns were quite strong when detected, we could not identify any differences in hand impairment, ipsilesional M1 excitability, or lesion location between stroke survivors in whom personalized patterns could and could not be detected. Although the sample size for these exploratory analyses was limited, these initial findings suggest that predicting whether personalized or group-level patterns will best reflect ipsilesional M1 excitability in a given participant is difficult. Identifying personalized EEG patterns also requires acquisition of large, exploratory TMS-EEG-EMG datasets in each participant, which is likely not feasible when delivering multi-day therapeutic interventions. Taken together, our findings identify bilateral sensorimotor mu rhythm suppression as a more feasible target for poststroke brain state-dependent TMS interventions than personalized EEG patterns.

The present study has its own limitations. First, we used post-hoc trial sorting to identify high and low excitability states in each stroke survivor. Future studies should validate our findings by delivering ipsilesional M1 TMS during periods of mu power suppression detected in real-time. Second, we used scalp-level EEG signals to identify power differences between excitability states. Although EEG provides excellent temporal resolution, its spatial resolution is limited. High-resolution structural MRI scans were unavailable for the vast majority of participants, preventing us from performing accurate source localization that accounted for each individual’s unique lesion characteristics. Third, the relationship between bilateral sensorimotor mu power suppression and ipsilesional M1 excitability was only present when evaluating high and low excitability states; linear regression did not reveal a significant effect of mu power on MEP amplitudes. This discrepancy suggests that mu power exerts its strongest effects on ipsilesional M1 excitability at extreme values. Given that the current study was performed at rest but that motor imagery, preparation, and execution all decrease mu power (Cassim et al., 2000; Duann & Chiou, 2016; Facchini et al., 2002; Pfurtscheller & Lopes da Silva, 1999; Szurhaj et al., 2003), future studies should confirm that volitional mu desynchronization further potentiates ipsilesional M1 excitability while also accounting for the influence of voluntary EMG activation.

In sum, this study is the first to systematically evaluate and characterize poststroke EEG brain states that distinguish between high and low ipsilesional M1 excitability. Our results challenge the notion that personalized EEG patterns can be used to index ipsilesional M1 excitability in all stroke survivors and instead identify bilateral mu power suppression as a novel and feasible target for therapeutic brain state-dependent TMS after stroke.

## Declaration of generative AI and AI-assisted technologies in the manuscript preparation process

During the preparation of this work, the author U Khatri used ChatGPT and Claude AI tools to assist with writing code for plotting and to refine the phrasing and clarity of the text at certain places. After using these tool/service, the author(s) reviewed and edited the content as needed and take(s) full responsibility for the content of the published article.

## Acknowledgments

Research reported in this publication was supported by the National Institute of Neurological Disorders as part of award # R21NSD133605, totaling $450,905 with 0% financed from non-federal sources. The content is solely the responsibility of the authors and does not necessarily represent the official views of the National Institutes of Health.

## Data availability

Raw data and analysis code are available upon reasonable request.

