## Supplementary Material for "Resting bilateral sensorimotor mu rhythm suppression facilitates ipsilesional M1 excitability after stroke"

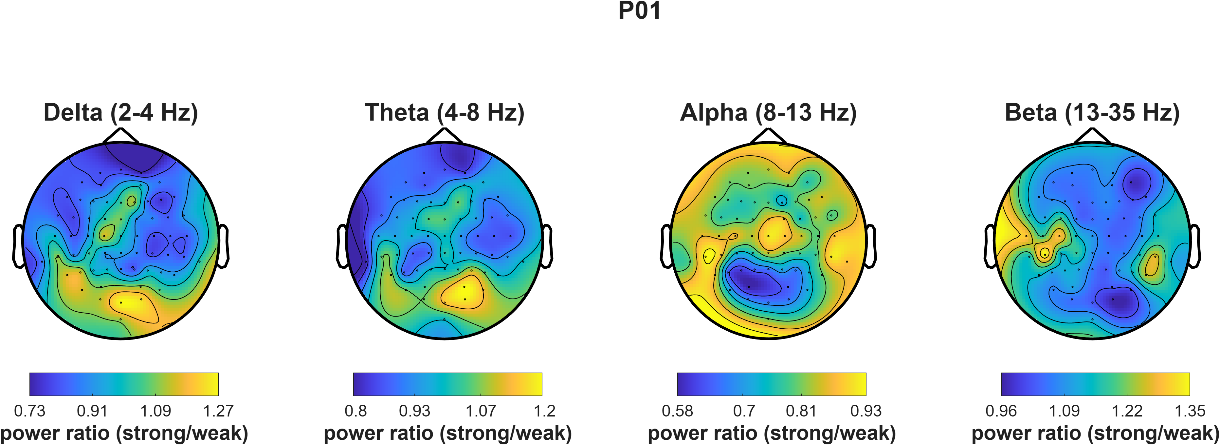

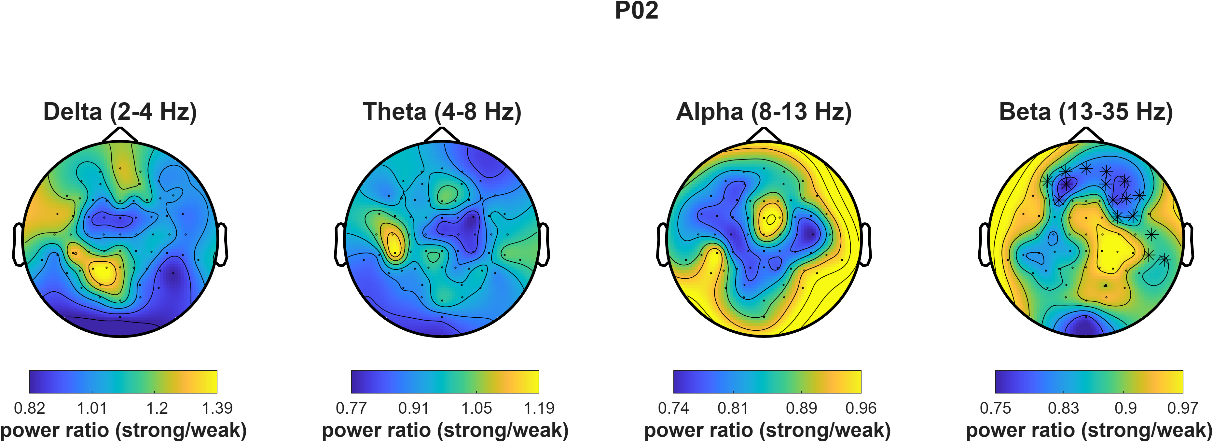

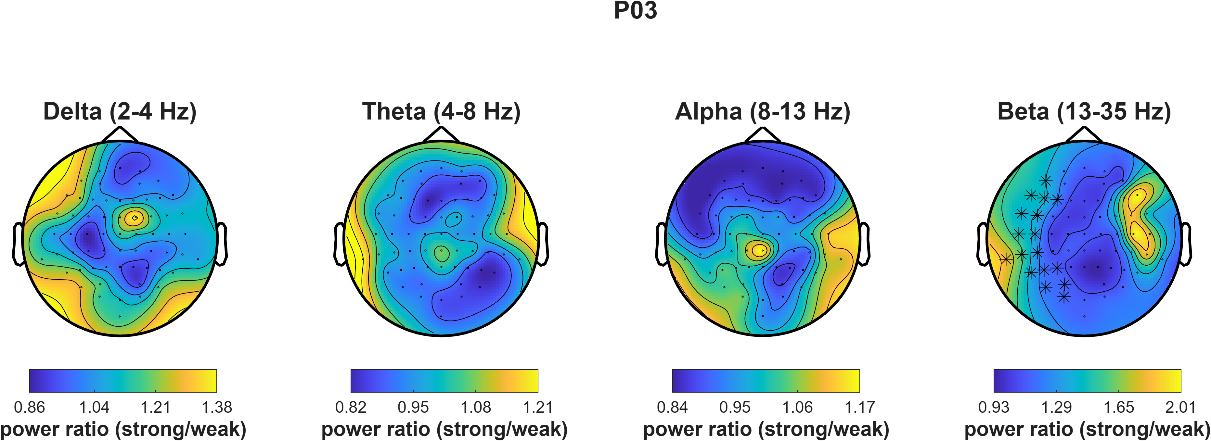

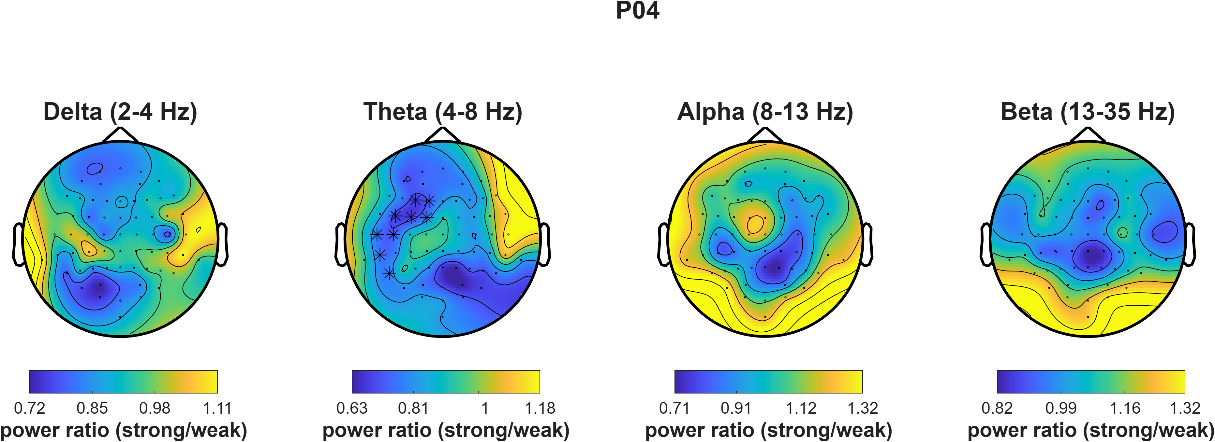

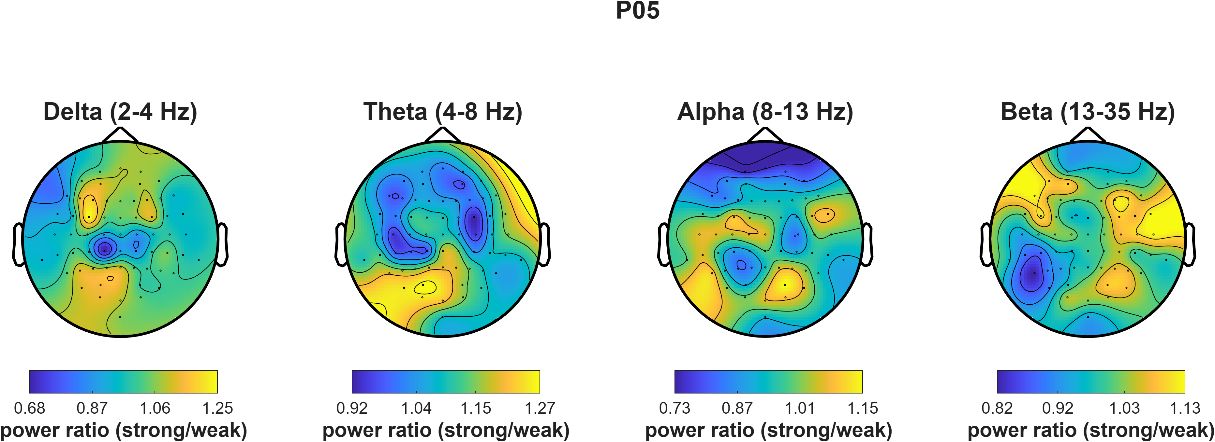

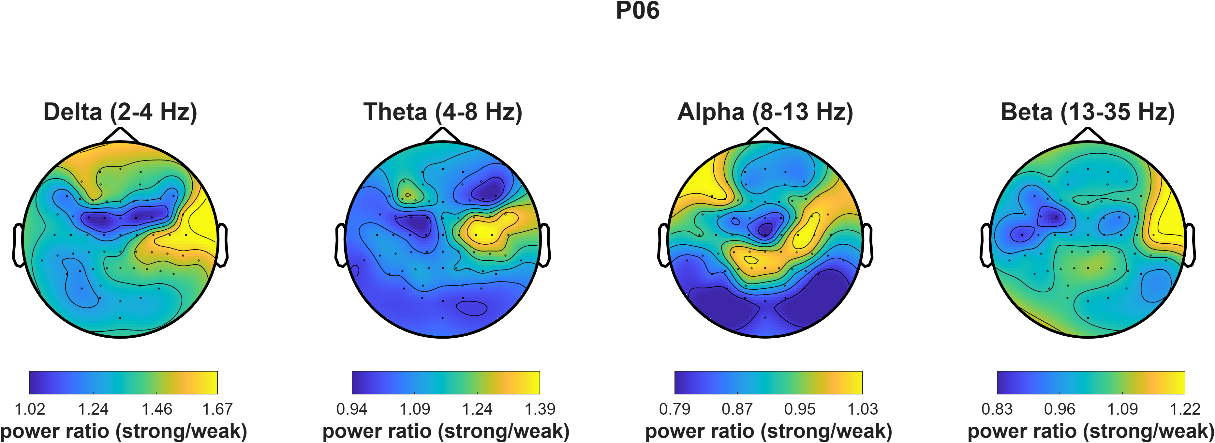

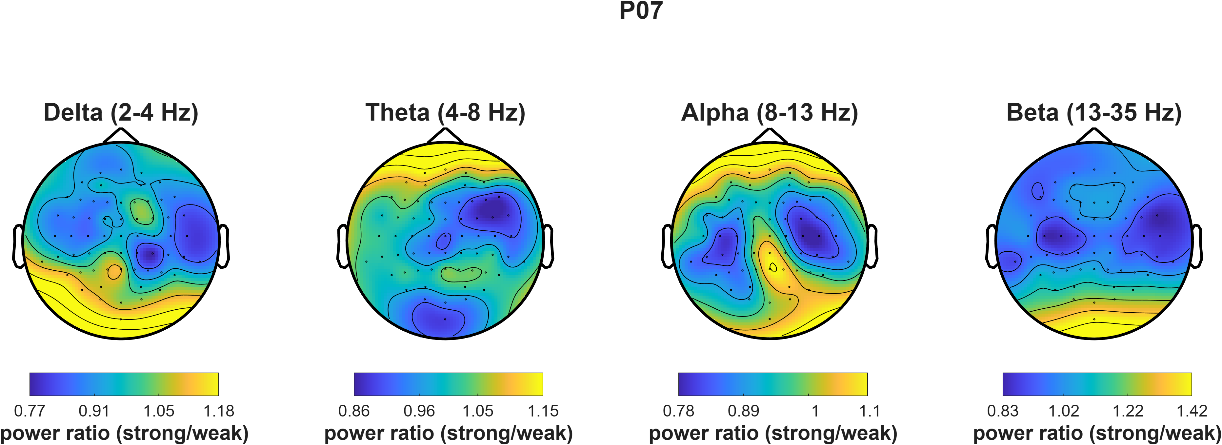

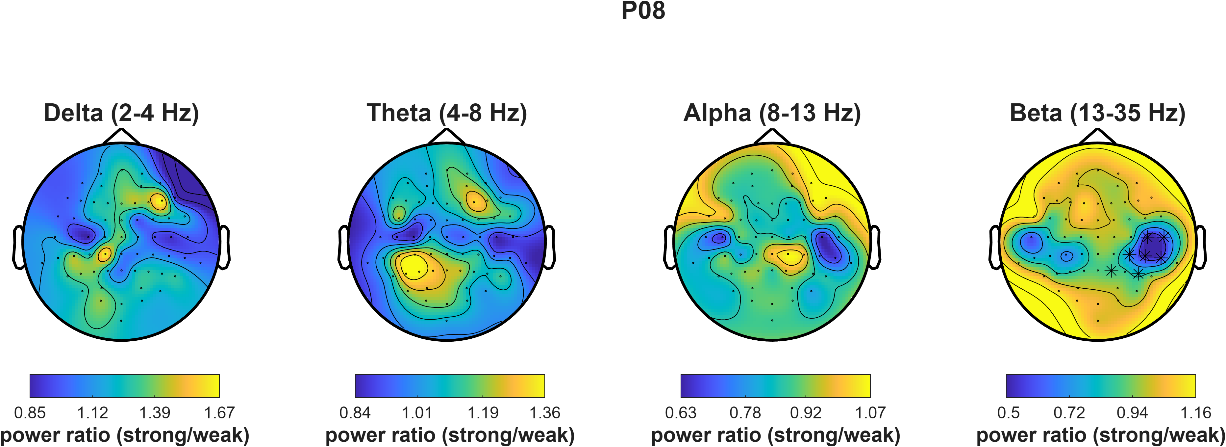

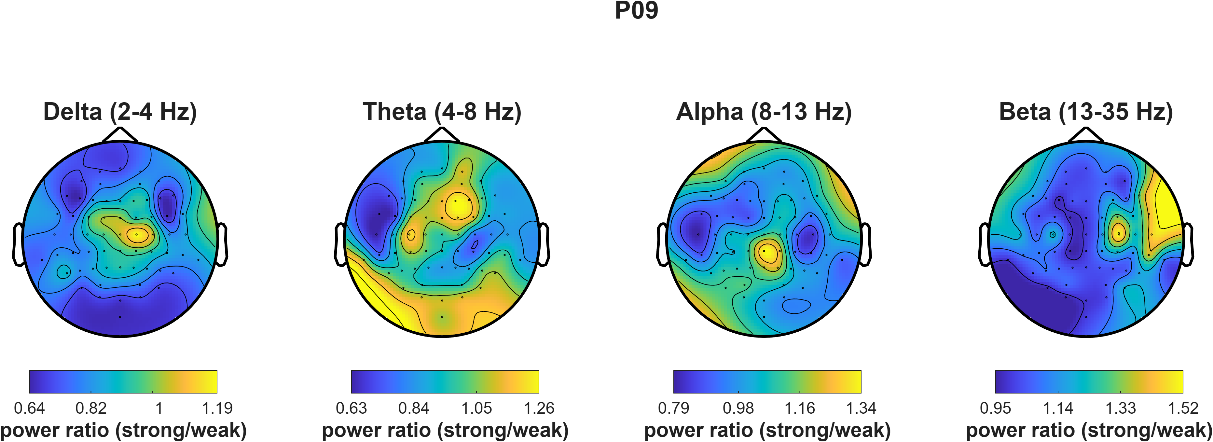

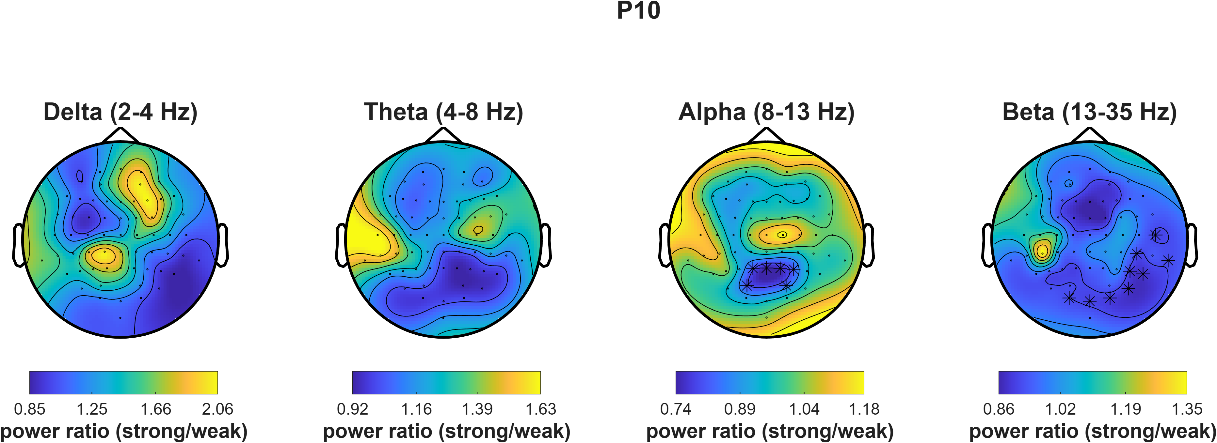

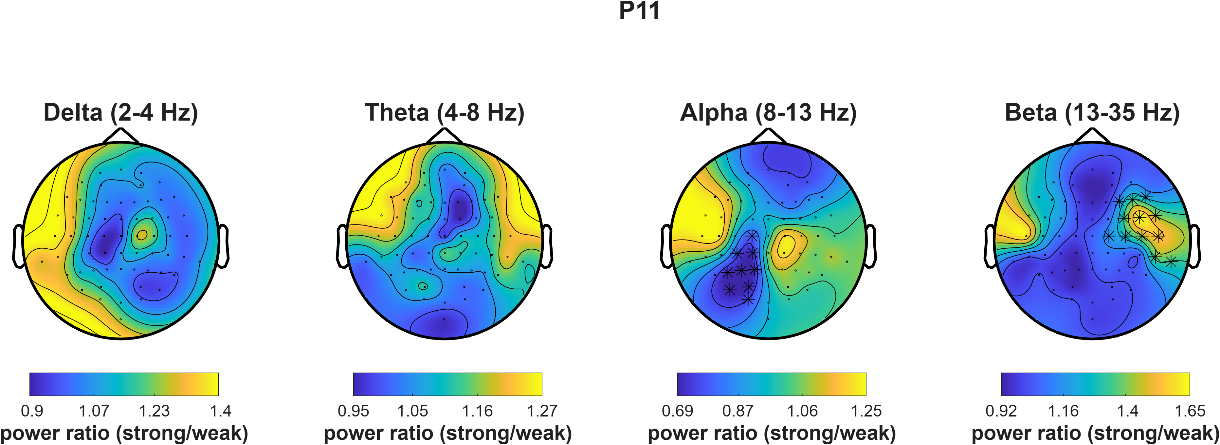

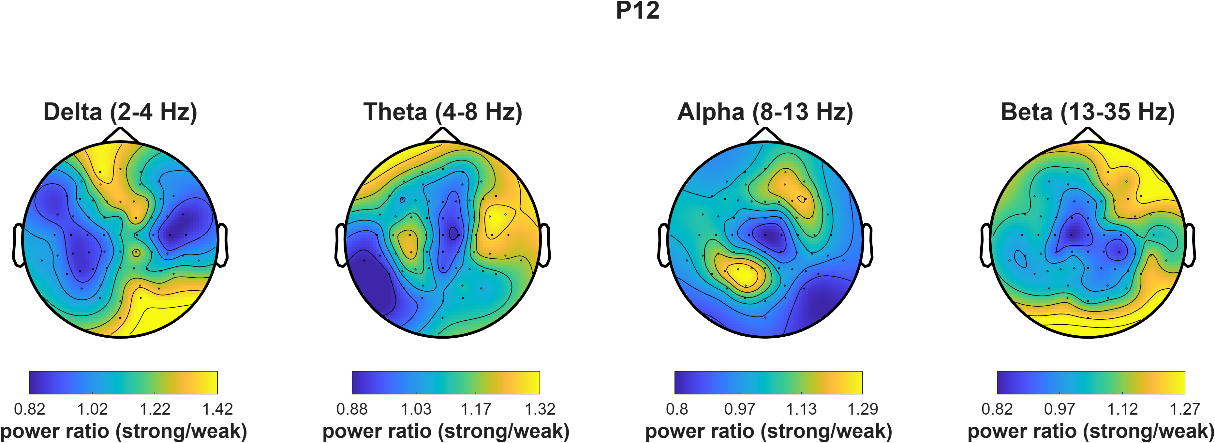

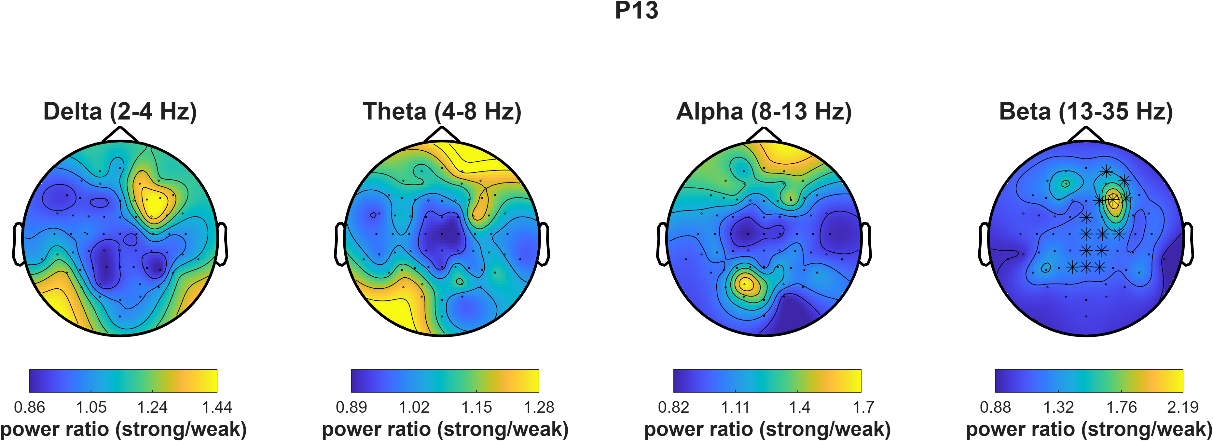

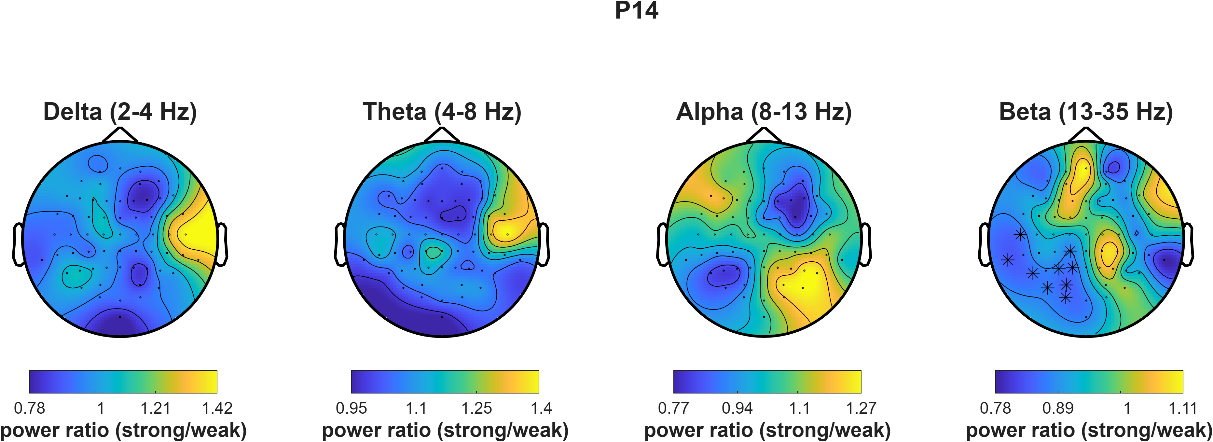

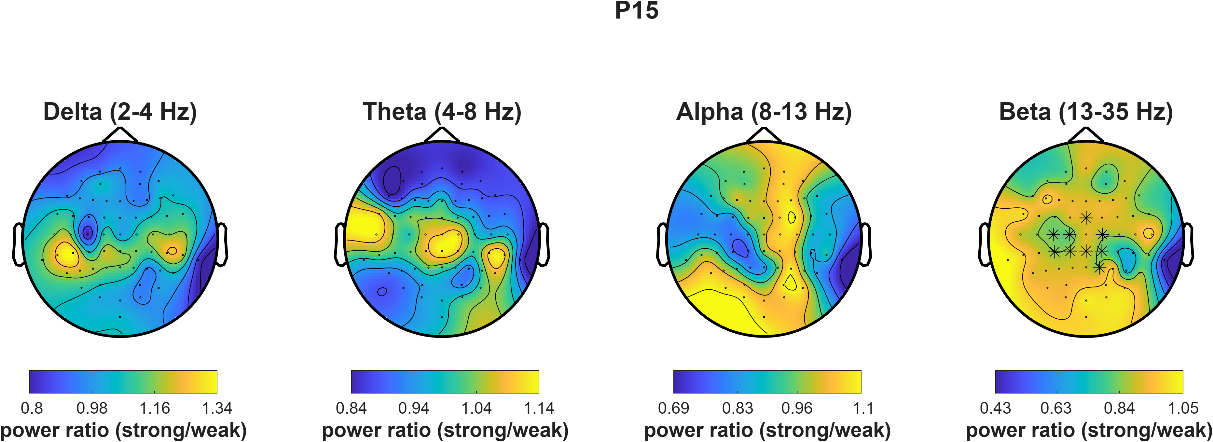

**Supplementary Figure 1. Subject-level EEG patterns distinguishing between high and low ipsilesional M1 excitability states.** Topoplots depicting EEG power ratios between high and low excitability states per frequency band. Asterisks and p-values indicate significant differences between excitability states obtained using cluster-based permutation tests. Xs indicate p < 0.05 and asterisks indicate p < 0.001.
